# A high-throughput fluorescence polarization assay to discover inhibitors of arenavirus and coronavirus exoribonucleases

**DOI:** 10.1101/2021.04.02.437736

**Authors:** Sergio Hernández, Mikael Feracci, Carolina Trajano De Jesus, Priscila El-Kazzi, Rafik Kaci, Laura Garlatti, Etienne Decroly, Bruno Canard, François Ferron, Karine Alvarez

## Abstract

Viral exoribonucleases are uncommon in the world of RNA viruses. To date, this activity has been identified only in the *Arenaviridae* and the *Coronaviridae* families. These exoribonucleases play important but different roles in both families: for mammarenaviruses the exoribonuclease is involved in the suppression of the host immune response whereas for coronaviruses, exoribonuclease is both involved in a proofreading mechanism ensuring the genetic stability of viral genomes and participating to evasion of the host innate immunity. Because of their key roles, they constitute attractive targets for drug development. Here we present a high-throughput assay using fluorescence polarization to assess the viral exoribonuclease activity and its inhibition. We validate the assay using three different viral enzymes from SARS-CoV-2, lymphocytic choriomeningitis and Machupo viruses. The method is sensitive, robust, amenable to miniaturization (384 well plates) and allowed us to validate the proof-of-concept of the assay by screening a small focused compounds library (23 metal chelators). We also determined the IC50 of one inhibitor common to the three viruses.

**Highlights:**

- *Arenaviridae* and *Coronaviridae* viral families share an exoribonuclease activity of common evolutionary origin
- *Arenaviridae* and *Coronaviridae* exoribonuclease is an attractive target for drug development
- We present a high-throughput assay in 384 well-plates for the screening of inhibitors using fluorescence polarization
- We validated the assay by screening of a focused library of 23 metal chelators against SARS-CoV-2, Lymphocytic Choriomeningitis virus and Machupo virus exoribonucleases
- We determined the IC_50_ by fluorescence polarization of one inhibitor common to the three viruses.

## 1. Introduction

RNA viruses code for a polyprotein, forming the replication/transcription complex (RTC) that orcherstrates viral replication. In this complex process, specific viral enzymatic activities are only found in some RNA viruses families. This is the case for the 3′-5′-exoribonuclease (ExoN) activity identified in *Arenaviridae* and *Coronaviridae* viral families, although these viruses display very different genome organization and replication/transcription strategies. Indeed, arenaviruses have an ambisense bi-segmented negative single-stranded RNA genome whereas the coronavirus genome is composed of a positive single-stranded RNA (Ferron et al., 2017; Knipe and Howley, 2013, Ogando et al., 2019). The ExoN activity identified in these two viral families are involved in different processes (Hastie et al., 2011; Snijder et al., 2003). Indeed, the ExoN carried by arenavirus nucleoprotein (NP) is likely involved in the suppression of the host innate immune response (Martínez-Sobrido et al., 2009, 2007, 2006) while the ExoN carried by the nsp14 protein of coronaviruses allows optimization of the replication fidelity of the viral genome (Bouvet et al., 2012; Ferron et al., 2018) as well as evasion of the host innate immunity (Becares et al., 2016; Lei et al., 2020).

The arenavirus and coronavirus ExoN belong to the same DEDD superfamily (DED/EDh subfamily), are structurally similar (Zuo and Deutscher, 2001) and are characterized by the presence of four conserved acidic residues (Asp and Glu) (Bouvet et al., 2012; Yekwa et al., 2019). The catalytic residues are essential for the binding of two metal cations involved in the RNA hydrolysis mechanism (Steitz and Steitz, 1993), always proceeding from the 3′- to the 5′-direction.

Because of its biological significance to the viral life cycle and role in infectious processes, arenavirus and coronavirus ExoN is an attractive target for drug development (Papageorgiou et al., 2020; Subissi et al., 2014). Currently, there are almost no FDA-approved antivirals against these viruses, despite the loss of human lives during the short-lived SARS-CoV outbreak in 2002 and the continuing MERS epidemic or regular outbreaks of LASSA virus in West Africa causing several thousand case fatalities. Today, the situation is different, the world is paralyzed by a global SARS-CoV-2 pandemic and this event revealed the urgency to act to develop potent antivirals, in support of a vaccine approach.

While millions of molecules are available in countless compounds libaries, drug development efforts are generally restricted by the limitations of validated therapeutic targets or available potent high-throughput screening assays. For mammarenavirus and coronavirus ExoN assay making use of radiolabeled substrate are available but barely adapted to HT screening (Bouvet et al., 2012; Ferron et al., 2018; Yekwa et al., 2019, 2017; Baddock et al., 2020; Saramago et al., 2021)).

Here we present a method using fluorescence polarization (FP) to assess the ExoN activity and its inhibition. The method relies on the hydrolysis of a fluorescent RNA substrate into smaller fragments which translates in a modification of the FP signal which is measured and recorded. The FP signal is altered proportionally to the size of the fluorescent RNA probe. We validate the method on three different viral enzymes of interest from SARS-CoV-2, Lymphocytic Choriomeningitis and Machupo viruses. The method is sensitive, robust, amenable to miniaturization (384 well plates) and allowed us to screen a focused library of 23 metal chelators over the three targeted viral ExoN, validating the proof-of-concept of the assay. We also determined the IC_50_ of one inhibitor common to the three viruses.

## 2. Materials and methods

### 2.1 Products and reagents

Raltegravir (**1**) and Baloxavir (**5**) were purchased from Carbosynth. Aurintricarboxyclic acid (**11**) was purchased from Acros Organics. Excepting for the compounds **8**, **13**, **14**, **15** and **24**, the other compounds have been described previously (Saez-Ayala et al., 2019, 2018). All compounds were resuspended in 100 % DMSO at 20 mM and stored at − 20 °C. The 22-mer RNA 5’-UGACGGCCCGGAAAACCGGGCC-3′ containing FAM dye at the 5′-end (5′-FAM-RNA) was purchased from Microsynth AG (Switzerland).

### 2.2 Protein expression and purification

The LCMV, MACV, ExoN plasmids used for this study were described in (Yekwa et al., 2019). The protein production method is detailed in the supplementary data file (see also figure S1 for elution profiles). For the expression of the SARS-CoV-2 nsp10 and nsp14, the synthetic genes were purchased from Twist (USA) and were cloned with a hexahistidine tag in the N-terminus of the protein. *E. coli* DE3 cells were transformed with the corresponding expression vectors. Bacteria were grown in TB medium with the corresponding antibiotic and protein expression was induced by addition of IPTG to a final concentration of 500 μM for nsp10 and 50 μM for nsp14 when the OD_600nm_ of the culture reached a value of 0.5. The induction was performed during 16 h at 17°C and 200 rpm. Bacterial cell pellets were frozen and resuspended in lysis buffer (50 mM HEPES, pH 7.5 and 300 mM NaCl) supplemented with 1 mM PMSF, 10 mM imidazole, 10 μg/ml DNase I, 0.25 mg/ml of lysozyme and 0.5% Triton X-100. After sonication and clarification, proteins were purified by two steps of chromatography. The first step consisted of an IMAC (Ni Resin). The lysate was passed through the column and washed with lysis buffer supplemented with 20 mM imidazole. The protein was eluted with lysis buffer supplemented with 250 mM imidazole. Protein fractions were then loaded on a HiLoad 16/60 Superdex 200 gel filtration column (GE Healthcare), and eluted with 10 mM HEPES, pH 7.5, 300 mM NaCl, 1 mM DTT and 5% glycerol. The fraction containing the pure protein, as examined by SDS-PAGE and coomasie staining, were pooled and concentrated in the gel filtration buffer, aliquoted in small volumes, flash frozen in liquid nitrogen and stored at −80°C.

### 2.3 Set-up of exonuclease activity conditions based on fluorescent polarization assay

#### 2.3.1 Optimisation of 5′-FAM-RNA concentration and SARS-CoV-2, LCMV or MACV ExoN ratio

Reactions were performed in 20 μl total volume in a buffer containing 40 mM Tris (pH 8), 5 mM DTT and 2 or 5 mM MnCl_2_. The concentration of 5′-FAM-RNA tested varied from 50 to 1000 nM. For each 5′-FAM-RNA concentration (50, 100, 250, 500 and 1000 nM), a range of concentration of ExoN (SARS-CoV-2, LCMV or MACV) was tested from a ratio of 5-fold less enzyme up to 10-fold more enzyme (10 nM to 10 uM) than 5′-FAM-RNA. For SARS-CoV-2 ExoN, the molar ratio of nsp14:nsp10 complex in the reactions was always kept at 1:4, as optimized previously (Bouvet et al., 2012). The reaction started by the addition of 5′-FAM-RNA and the fluorescence polarization (FP) was read in the Pherastar FSX (BMG Labtech) using the 480 nm excitation and 520 nm emission filter, during 30 minutes at 25°C, every 30 seconds. The gain was set up using the negative control which contained the fluorescent RNA in the reaction buffer with metal ion and in the presence of heat-denatured nuclease. Other negative controls tested used consisted in the replacement of MnCl_2_ by CaCl_2_ or, depletion of the metal ion in the reaction mix and depletion of the enzyme. After 30 minutes, the reactions were stopped by the addition of an equal volume of loading buffer (8M urea containing 10 mM EDTA) and the digestion products were then loaded in 7 M urea containing 20% (wt/vol) polyacrylamide gels (acrylamide/bisacrylamide ratio 19:1) buffered with TBE and visualized using a Fluorescent Image Analyzer Typhoon (GE Healthcare).

#### 2.3.2 Optimisation of temperature, metal ion nature and concentration requirement

To find the optimal temperature, metal ion nature and reagent concentration, reactions were performed in the buffer mentioned above. The 5′-FAM-RNA concentration was kept at 100 nM and ExoN concentration ranged from 100 nM to 1600 nM. The activity was tested in the presence of MnCl_2_ or MgCl_2_ at three different concentrations 1, 2 or 5 mM. The FP signal was recorded as mentioned previously either at 25°C or at 37°C. After finishing the recording of the FP signal, an aliquot of the sample was examined in urea PAGE as mentioned above.

#### 2.3.3 Time course of ExoN assay

Reactions of 20 μl total volume were performed in the buffer mentioned above, with 2 mM MnCl2 for SARS-CoV-2 ExoN and 5 mM for LCMV or MACV ExoN. The 5’-FAM-RNA concentration was fixed at 100 nM. The SARS-CoV-2 nsp14/nsp10 ExoN complex concentration was ranged from 200 nM to 1 μM and pre-incubation at RT during 5 minutes was performed to allow the complex formation. The LCMV and MACV ExoN concentration was ranged from 100 nM to 1600 nM. The reaction started by the addition of 5′-FAM-RNA and The FP signal was recorded as mentioned previously. After finishing the recording of the FP signal, an aliquot of the sample was examined in urea PAGE as mentioned above. The assay was done in triplicate for each ExoN.

### 2.4 High throughput screening and IC_50_ determination

#### 2.4.1 HTS assay based on fluorescent polarization assay

The screening at 5 μM and 20 μM inhibitor concentration was performed in 384-wells with flat bottom Nunc plates, in 20 μL total volume reaction. Reactions were performed in a buffer containing 40 mM Tris (pH 8), 5 mM DTT, with 2 mM MnCl_2_ for SARS-CoV-2 and 5 mM MnCl_2_ for LCMV or MACV ExoN. The 5′-FAM-RNA concentration was fixed at 100 nM. The ExoN concentration was fixed at 400 nM. Compounds (**1**-**23**) were added to the reaction with final concentration of 5 μM and 20 μM in 5% DMSO. Reactions were initiated by addition of the 5′-FAM-RNA using a multichannel pipette. 48 positive controls (the reaction mix with 5% DMSO, with ExoN and without compound) have been deposited randomly in the 384-well plate. 48 negative controls (the reaction mixture with 5% DMSO, without or with ExoN that has been heat denatured, and without compound), have been deposited randomly in the 384-well plate. The reaction started by the addition of 5′-FAM-RNA and the FP signal was recorded as mentioned previously. To calculate the percentage of inhibition, a time correction was applied for the delayed initiation of the reaction due to the use of a multichannel pipette. The percentage of inhibition at a given time was calculated as follows:

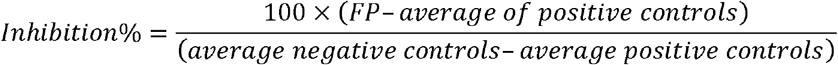

where FP correspond to the fluorescent polarization signal of a compound.

The time selected for doing this calculation was the time when the signal of the positive control reached the plateau (30 minutes). The Z’ factor for the assay was calculated using the following equation:

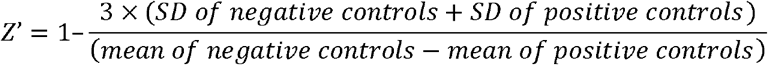

where SD is the standard deviation. 48 positive controls and 48 negative controls were considered to calculate the Z’ factor.

#### 2.4.2 IC_50_ determination of compound 11 based on fluorescent polarization assay

Reactions were performed in a buffer containing 40 mM Tris (pH 8), 5 mM DTT, with 2 mM MnCl_2_ for SARS-CoV-2 ExoN and 5 mM for LCMV or MACV ExoN. The 5′-FAM-RNA concentration was fixed at 100 nM. The ExoN concentration was fixed at 400 nM. The compound **11** concentration varied from 0.2 to 12.5 μM for LCMV ExoN, from 0.4 to 25 μM for MACV ExoN and from 0.5 to 16 μM for SARS-CoV-2 ExoN nsp14/nsp10 complex. The reaction started by the addition of 5′-FAM-RNA and the FP signal was recorded as mentioned previously. The percentage of inhibition was calculated as indicated in the previous section. The curves of percentage of inhibition respect to the inhibitor concentration in a logarithmic scale were fitted in Graphpad Prism software using a four parameters equation. The assays were done in triplicate.

## 3. Results and discussion

Fluorescence polarization (FP) is a reliable and sensitive tool for monitoring enzymatic reaction progress, by determining the difference of polarization signals during reaction (Zhang et al., 2012). FP - based assays in HTS studies have been used in drug discovery (Lea and Simeonov, 2011; Uri and Nonga, 2020, Baughman et al., 2012; Liu et al., 2014)). The FP signal recorded is proportional to the molecular weight (MW) of a fluorescent molecule (Kwok, 2002; Latif et al., 2001). We decided to apply this method to viral ExoN activity, by monitoring the size of a fluorescent labeled RNA probe which is altered in the course of the nuclease activity, reflecting the enzymatic activity (Liu et al., 2014; Zhang et al., 2012). Because several factors can change the FP- monitoring success, the assay was first optimized.

### 3.1 Determination of optimized experimental conditions of the SARS-CoV-2 nsp14/nsp10 complex, LCMV and MACV ExoN activity based on FP assay

To set-up a HTS assay for the SARS-CoV-2, LCMV and MACV ExoN activity, we explored 5’-FAM-RNA substrate(s) and enzyme(s) concentration, metal ion co-factors, temperature and reaction duration. RNA substrate and optimal conditions for arenavirus and coronavirus ExoN activity are already described (Bouvet et al., 2012; Saramago et al., 2021; Yekwa et al., 2019, 2017). A 22mer RNA that forms stable hairpin in its 3’ end has been reported to be a valuable substrate for both arenavirus and coronavirus ExoN. Moreover, for SARS-CoV-2 nsp14 ExoN, nsp10 was added in the reaction (ratio 1:4 of nsp14:nsp10) as nsp10 was previously demonstrated to stimulate > 35-fold the nsp14 ExoN activity (Bouvet et al., 2012; Saramago et al., 2021).

We first determined the smallest 5′-FAM-RNA concentration that provides sufficient FP signal, concomitantly with the set-up of the ratio of ExoN and 5′-FAM-RNA to obtain a reproducible and stable FP signal. For each 5′-FAM-RNA concentration tested (50 to 1000 nM), single turn over (STO) conditions (excess of enzyme respect to substrate) and also multiple turn over (MTO) conditions (excess of substrate respect to enzyme) were tested. Because the RNA substrate is degraded during the ExoN activity, FP value is altered proportionally. A significant change in the FP signal due to the hydrolysis of the 5′-FAM-RNA was observed only under STO conditions. 100 nM concentration of 5′-FAM-RNA was selected because it was the smallest concentration of substrate providing a reproducible and stable FP signal.

We optimized the positive and negative controls of ExoN activity. Several combinations were tested (Supplementary figure S2): 5′-FAM-RNA in reaction buffer with metal ions, 5′-FAM-RNA in reaction buffer with ExoN and without metal ions, 5′-FAM-RNA in reaction buffer with ions and denaturated ExoN and 5′-FAM-RNA in reaction buffer with CaCl_2_ instead of MnCl_2_. For all the negative controls tested, there was no variation between the initial and final FP values after 30 min incubation, indicating the absence of degradation of the fluorescent probe. However, we observe that the initial FP values are lower without metal ions in the reaction buffer (Supplementary figure S2), or in the presence of EDTA 10 mM (data not shown). This results is in agreement with the study of Liu *et al* (Liu et al., 2014), who carried out the experiments to investigate the influence of metal ions in FP assays, and reported that cations concentration can affect the FP signal by altering the mobility of the fluorophore through stabilizing the RNA secondary structure. Thus, for the rest of the study, the negative controls were prepared by mixing 5′-FAM-RNA in reaction buffer with ions and without ExoN.

Regarding the nature of the metal ion, MnCl_2_ and MgCl_2_ were tested. An increased activity is observed in the presence of MnCl2 for both arenaviruses and SARS-CoV-2 ExoN (data not shown). The concentration of MnCl_2_ that shows the highest reduction in FP signal is 5 mM for arenaviruses as previously reported (Yekwa et al., 2019, 2017) and 2 mM for SARS-CoV-2 as recently reported (Baddock et al., 2020; Saramago et al., 2021). Finally the temperature selected to perform the assay is 25°C since the incubation at 37°C leads to a significative evaporation of the sample that reduces the stability and reproducibility of the FP signal.

### 3.2 Assessing the SARS-CoV-2, LCMV and MACV ExoN activity by FP

While the 5’-FAM-RNA substrate was fixed at 100 nM, which was the lowest concentration fairly detected by the sensitivity of the method, different concentrations of ExoN were tested (100 nM to 1600 nM). The FP signal change was measured during 30 min. The FP curves and digestion products examined in urea PAGE after FP reading, are gathered in figure 1.

**Figure 1.**
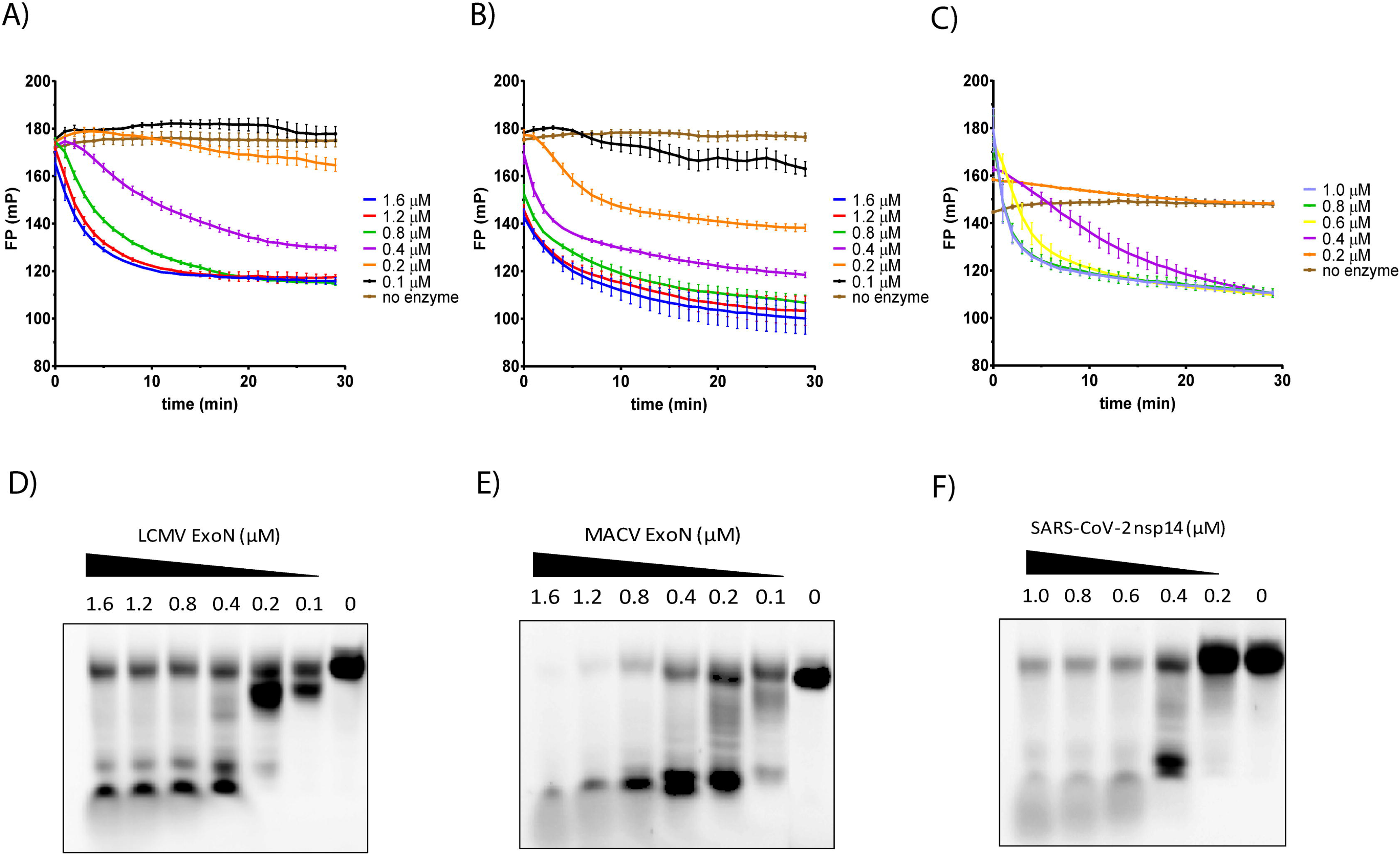
ExoN activity measured by FP for LCMV (panel A), MACV (panel B) and SARS-CoV-2 (panel C). The FP signal variation is recorded with time during 30 min, each 30 secondes, at 25°C. The 5′-FAM- RNA substrate concentration used is 100 nM and the ExoN concentration tested ranges 100 nM to 1600 nM. The data represents the average and SEM of three independent experiments. The bottom panels illustrate a representative image of the digestion products analyzed in urea PAGE, of the ExoN activity for LCMV (panel D), MACV (panel E) and SARS-CoV-2 (panel F), after recording the FP signal. An aliquot of the sample was loaded into a urea-PAGE 20% and scanned in a fluorescence imager.

The negative controls (figure 1, brown curves, in panels A, B and C) present flattened curves, as expected. Remarkably, early in the reaction a small increase in the initial FP value is observed, which could correspond to the time required for the formation of the catalytic complex between the ExoN, the catalytic cations and the 5′-FAM-RNA substrate. This is more visible for the SARS-CoV-2 ExoN, which requires the interaction of nsp14 and nsp10 forming the active complex. This increase of fluorescence occurs for all FP curves but is visible mainly on the FP curves corresponding to the lowest ExoN concentrations when the formation of the pre-hydrolytic complex presumably takes the longest time.

For all tested ExoN, excepting for the lowest used concentration, we observe a reduction of the FP signal with time. The decrease of the FP value is correlated with the hydrolysis of the 5’-FAM-RNA substrate by the ExoN, as confirmed on the urea PAGE images (figure 1, panels D, E and F). The difference between initial and final FP value after 30 min is proportional to the number of nucleotides removed from the 5′-FAM-RNA.

By increasing the ExoN concentration, we observe an increase in the FP curves slope but the FP signals reach a plateau which might be related to the inability to remove any extra nucleotide after a certain point.

The method efficiency was validated by the difference of FP values between the negative control (figure 1, brown curves, in panels A, B and C) and the FP value obtained at one concentration after 30 min reaction. This difference was selected as significant enough, with ΔFP +/- 30 units of mP.

To validate this method and use it as screening assay, we decided to use a molar ratio 1:4 between the 5′-FAM-RNA substrate (100 nM) and ExoN (400 nM). With the optimized duration, temperature and experimental conditions, this ratio gives the more stable and reproducible FP signals, correlated with specific 5’-FAM-RNA fragments.

### 3.3 Screening of a focused library against the SARS-CoV-2, LCMV and MACV ExoN using FP

Because ExoN activity is metal dependent, we tested a focused library of 23 metal chelators that we have developed previously (Saez-Ayala et al., 2019, 2018), in order to demonstrate the robustness of our FP method. Two screenings were performed using 5 μM and 20 μM of compounds in 5% DMSO in a single assay. The robustness of the assay for HTS was calculated using 48 negative and 48 positive controls. The lowest Z′ value is 0.68 indicating that the assay is reliable (see supplementary data figure S3). The percentages of inhibition at 5 μM and 20 μM for each ExoN, extracted from FP curves (see supplementary data figure S4), are gathered in supplementary figure S5 and figure 2 (panels A, B and C), respectively. The degree of digestion of the RNA substrate was also controlled by fragments separation on urea-PAGE at the end of the reaction for compounds **4**, **11**, **13**, **18** and **23**(figure 2, panel D).

**Figure 2.**
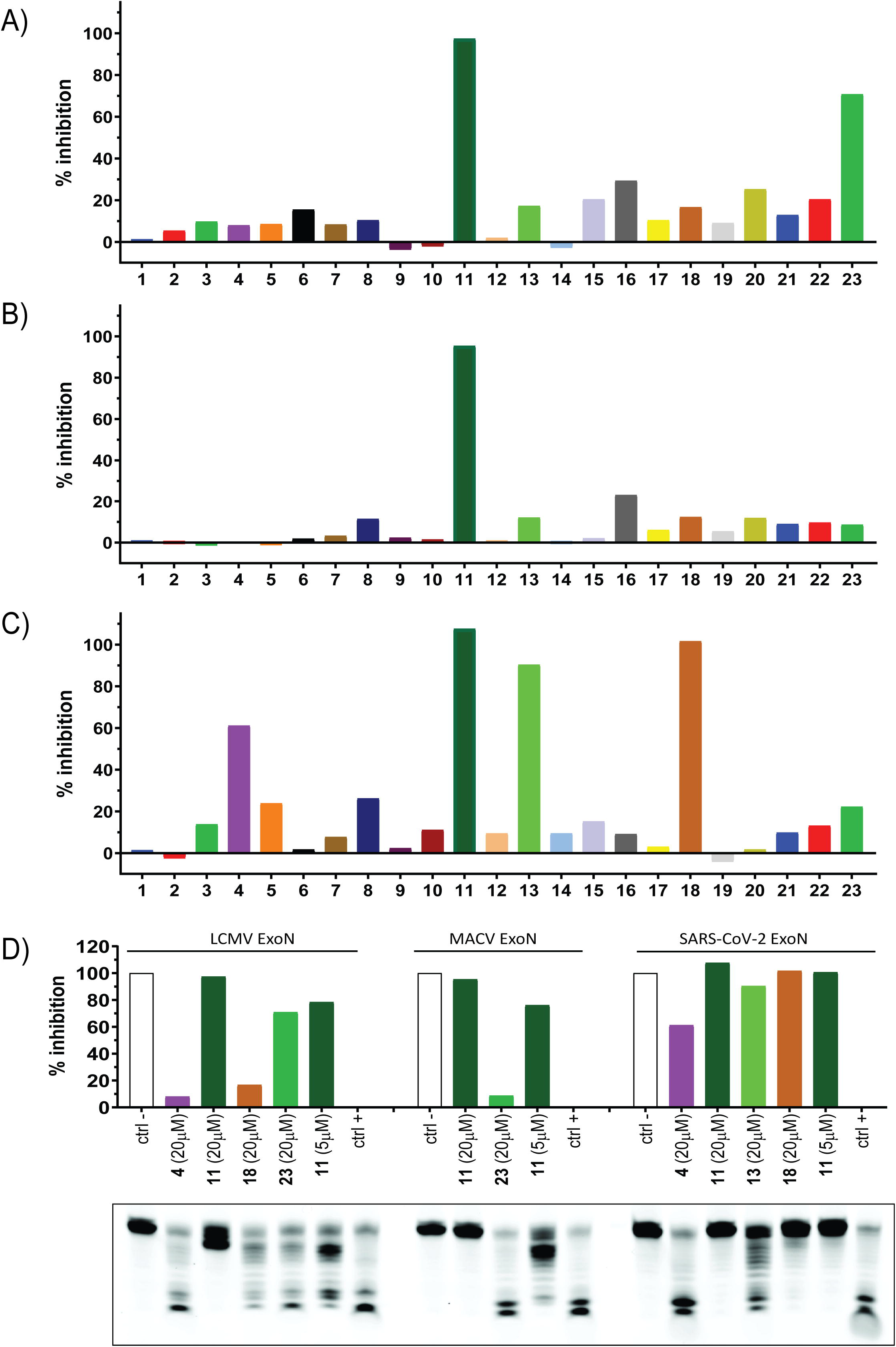
Screening of focused library of metal chelators (23) against LCMV (panel A), MACV (panel B) and SARS-CoV-2 (panel C) ExoN followed by FP. Orthogonal validation assay by analysis of the fluorescent RNA substrate by urea PAGE (panel D) The bars show the % of inhibition of the ExoN activity as described in materials and methods. For the screening conditions 100 nM 5′-FAM-RNA, 400 nM ExoN and 20 μM of inhibitor were used.

We identified compound **11** which inhibits 100% of the different ExoN activity at 20 μM and 78%, 76% and 100% at 5 μM, respectively against LCMV, MACV and SARS-CoV-2 ExoN. Compound **11** which is Aurintricarboxyclic Acid (ATA) was included as positive control as it has been previously described as nuclease inhibitor (Huang et al., 2016), acting as non-specific metal chelator (Sharma et al., 2000). We also identified inhibitors showing more specific inhibition profile. Compound **4** is active against SARS-CoV-2 with 61% inhibition at 20 μM (less than 10% inhibition of LCMV ExoN) while compound **23** is specific to LCMV ExoN with 71% inhibition (less than 10% inhibition of MACV ExoN). Compounds **13** and **18** are potent inhibitors of SARS-CoV-2 ExoN with more 90% and weak inhibitors of LCMV and MACV ExoN (less than 20% inhibition). The FP method allowed the identification of potent ExoN inhibitors as compounds **4, 11**, **13**, **18** and **23**, and the absence of RNA degradation was confirmed by urea PAGE analysis (figure 2 - panel D) used as an orthogonal validation assay.

### 3.4 Assessing IC_50_ measurement of compound **11** by FP

Compound **11** displaying the highest inhibition, at both 5 and 20 μM, was used as model for IC_50_ determination. The FP curves, the digestion products examined in urea PAGE after FP reading and dose-response curves obtained by extraction of FP data are gathered in figure 3. The IC_50_ values were determined by hill plot curve fitting (see materials and methods section). The IC_50_ values for LCMV, MACV and SARS-CoV-2 ExoN were 3.61 ± 0.20 μM, 3.46 ± 0.29 μM and 1.64 ± 0.17μM, respectively. Notably, these values correlated perfectly with the 80% inhibition obtained by screening at 5 μM, confirming the robustness and reproducibility of the assay. The FP signal correlates also with the degree of digestion observed by gel.

**Figure 3.**
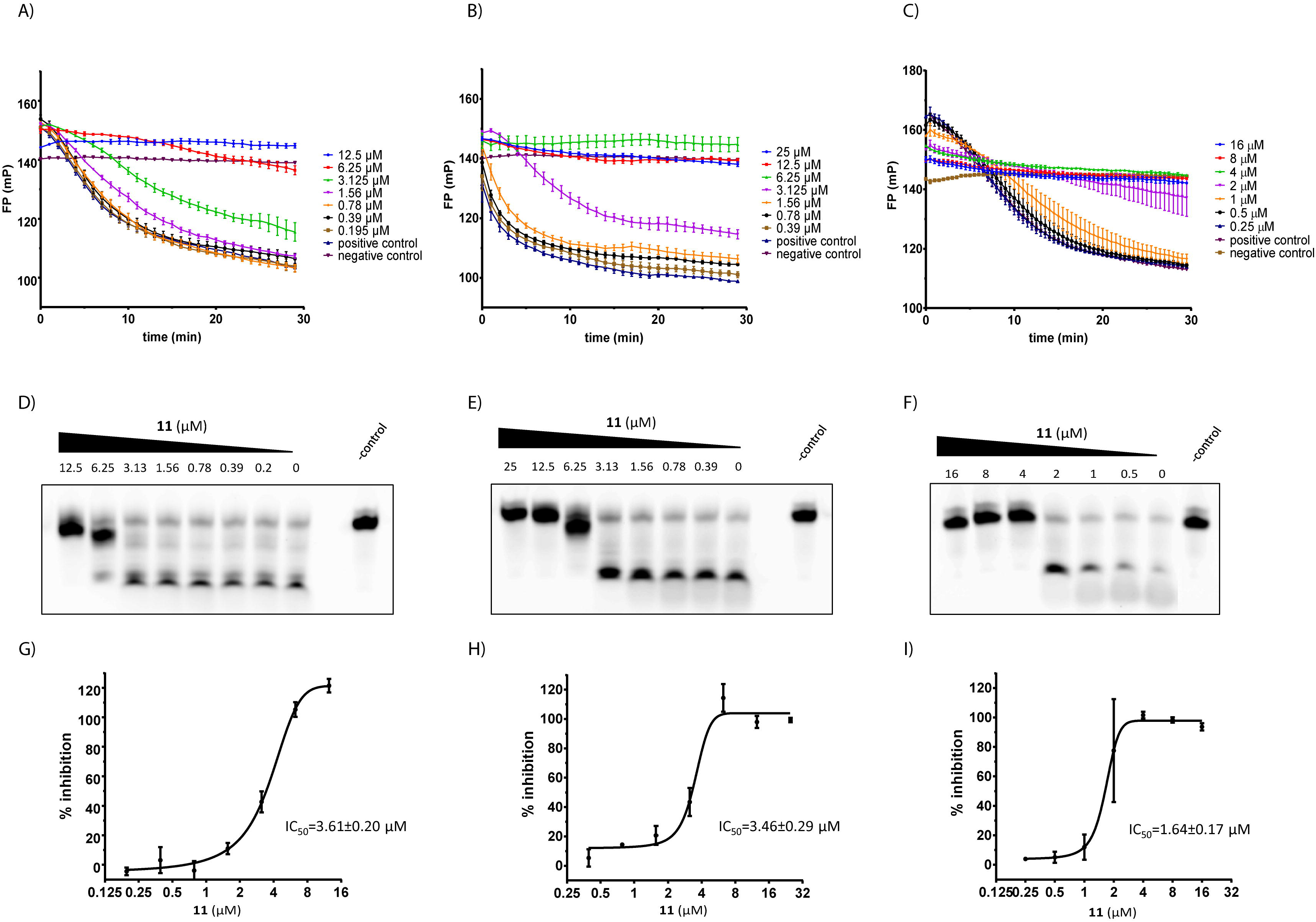
IC_50_ measurement of compound **11** by FP on LCMV, MACV and SARS-CoV-2 ExoN. In A, B and C the FP signal variation is recorded with time during 30 min, each 30 seconds, at 25°C. The 5′-FAM-RNA substrate and LCMV (panel A), MACV (panel B) and SARS-CoV-2 (panel C) ExoN concentration were respectively 100 nM and 400 nM. The data represents the average and SEM of three independent experiments. In the middle panels it is shown a representative image of the digestion products of the LCMV (panel D), MACV (panel E) and SARS-CoV-2 (panel F) ExoN in presence of the different concentrations of compound **11**, visualized in urea AGE after recording the FP signal. In the bottom panels it is shown the dose-response curves obtained by extraction of FP data, the IC_50_ values for compound **11** with LCMV (panel G), MACV (panel H) and SARS-CoV-2 (panel I). The data represents the average and SEM of three independent experiments.

## 4. Conclusion

The use of fluorescence polarization assay for inhibitors identification was investigated and we prove that this technique is reliable and sensitive to monitor nuclease activity.

Our work presents the development of a viral ExoN HTS assay in 384-well plates. Its most valuable feature is that allows a reliable and rapid identification of ExoN inhibitors limiting the ExoN activity of viral enzymes belonging to arenavirus (LCMV and MACV) and coronavirus (SARS-CoV-2).

These results make fluorescence polarization assay an important tool in the screening of compounds libraries to discover antivirals.

## Funding and Acknowledgments

This work was supported by grants from the Ministry of the Armed Forces (DGA) – Defense Innovation Agency (AID) and the French National Research Agency (ANR-18-ASTR-0010-01, PaNuVi), Fondation pour la Recherche Medicale (Chemistry for Medecine, DCM20181039531), SCORE project H2020 SC1-PHE Coronavirus-2020 (grant# 101003627). Priscila El-Kazzi and Rafik Kaci were funded by Fondation Mediterranée Infection (Infectiopole Sud), Laura Garlatti was funded by DGA and Aix-Marseille University (fellowship N° DGA01D19024292 AID). The Carolina Trajano De Jesus master’s internship was supported by the Coordenação de Aperfeiçoamento de Pessoal de Nível Superior - Brasil (CAPES) – Finance Code 001. We thank Thi Hong Van Nguyen and Adrien Delpal for technical assistance.

Supplementary data associated with this article can be found, in the online version, at

