## Supplementary data file for "A high-throughput fluorescence polarization assay to discover inhibitors of arenavirus and coronavirus exoribonucleases"

**Supplementary methods:**

***Protein expression and purification***

For the expression of LCMV and MACV exonucleases, the genes were cloned in the pETG20A vector (Invitrogen) which adds a cleavable thioredoxin-hexahistidine tag in the *N*-terminus of the protein. E. coli DE3 cells were transformed with the corresponding expression vectors. Bacteria were grown in TB medium with the corresponding antibiotic and protein expression was induced by addition of IPTG to a final concentration of 500 µM and 100 µM ZnCl_2_ when the OD600nm of the culture reached a value of 0.5. The induction was performed during 16 h at 17°C and 200 rpm. Bacterial cell pellets were frozen and resuspended in lysis buffer (50 mM HEPES, pH 7.5 and 300 mM NaCl) supplemented with 1 mM PMSF, 10 mM imidazole, 10 µg/ml DNase I, 0.25 mg/ml of lysozyme and 0.5% Triton X-100. After sonication and clarification, proteins were purified by two steps of chromatography. The first step consisted of an IMAC (Ni Resin). The lysate was passed through the Nickel resin and washed with lysis buffer supplemented with 20 mM imidazole. The protein was eluted with lysis buffer supplemented with 250 mM imidazole. The thioredoxin-hexahistidine tag was later removed by cleavage with TEV protease followed by a second nickel affinity chromatography. Untagged protein fractions were then loaded on a HiLoad 16/60 Superdex 75 gel filtration column (GE Healthcare), and eluted with 20 mM HEPES, pH 7.5, 300 mM NaCl and 5% glycerol. The fraction containing the pure protein, as examined by SDS-PAGE and coomasie staining, were pooled and concentrated in the gel filtration buffer, aliquoted in small volumes, flash frozen in liquid nitrogen and stored at -80°C.

For the expression of the SARS-CoV-2 nsp10 and nsp14, the genes were cloned with a hexahistidine tag in the N-terminus of the protein. E. coli DE3 cells were transformed with the corresponding expression vectors. Bacteria were grown in TB medium with the corresponding antibiotic and protein expression was induced by addition of IPTG to a final concentration of 500 µM for nsp10 and 50 µM for nsp14 when the OD600nm of the culture reached a value of 0.5. The induction was performed during 16 h at 17°C and 200 rpm. Bacterial cell pellets were frozen and resuspended in lysis buffer (50 mM HEPES, pH 7.5 and 300 mM NaCl) supplemented with 1 mM PMSF, 10 mM imidazole, 10 µg/ml DNase I, 0.25 mg/ml of lysozyme and 0.5% Triton X-100. After sonication and clarification, proteins were purified by two steps of chromatography. The first step consisted of an IMAC (Ni Resin). The lysate was passed through the Nickel resin and washed with lysis buffer supplemented with 20 mM imidazole. The protein was eluted with lysis buffer supplemented with 250 mM imidazole. Protein fractions were then loaded on a HiLoad 16/60 Superdex 200 gel filtration column (GE Healthcare), and eluted with 10 mM HEPES, pH 7.5, 300 mM NaCl, 1 mM DTT and 5% glycerol. The fraction containing the pure protein, as examined by SDS-PAGE and coomasie staining, were pooled and concentrated in the gel filtration buffer, aliquoted in small volumes, flash frozen in liquid nitrogen and stored at -80°C.

**Supplementary figures.**

***Supplementary figure S1*.** Purification of viral ExoN. Elution profile after gel filtration in superdex S75 16/600 and SDS-PAGE and coomasie stain of proteins after final step of purification for A) SARS-CoV-2, B) LCMV and C) MACV ExoN. The peak corresponding to the MACV ExoN is highlighted in the red box.

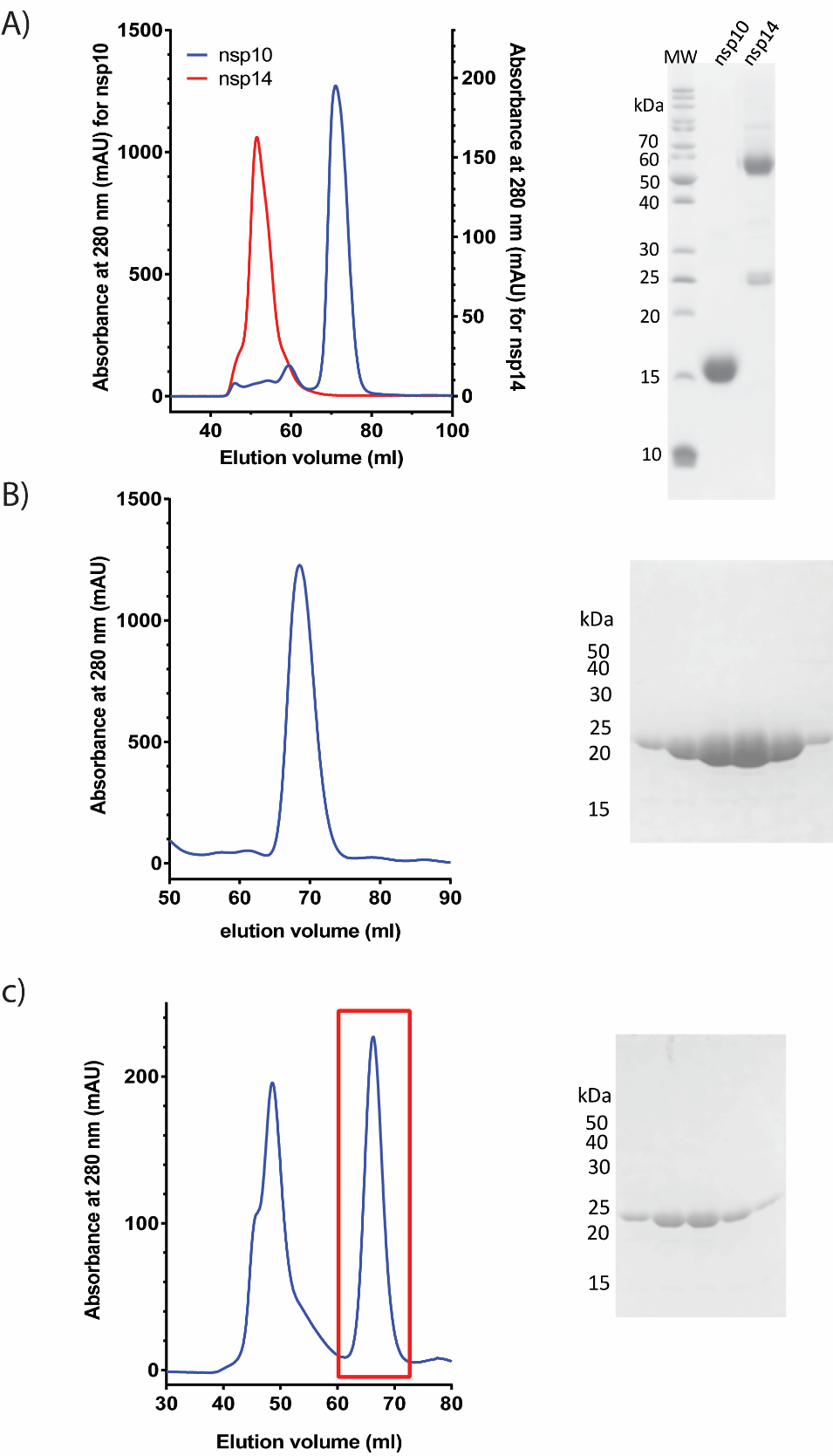

***Supplementary figure S2.*** A) Fluorescence polarization profiles of different negative controls and positive controls using the LCMV ExoN. The exoribonuclease reaction was performed as described in material and methods. The positive control contains the reaction mixture in the presence of the exoribonuclease, the CaCl_2_ consist in the full reaction mixture where the MnCl_2_ has been replaced by CaCl_2_, a negative control was also included where the enzyme was heat inactivated (denatured enzyme), a control lacking metal ions also was included, and finally a control in presence of the reaction mix lacking the ExoN. B) urea PAGE showing the degradation profile of the different controls mentioned above after 30 min incubation at 25°C.

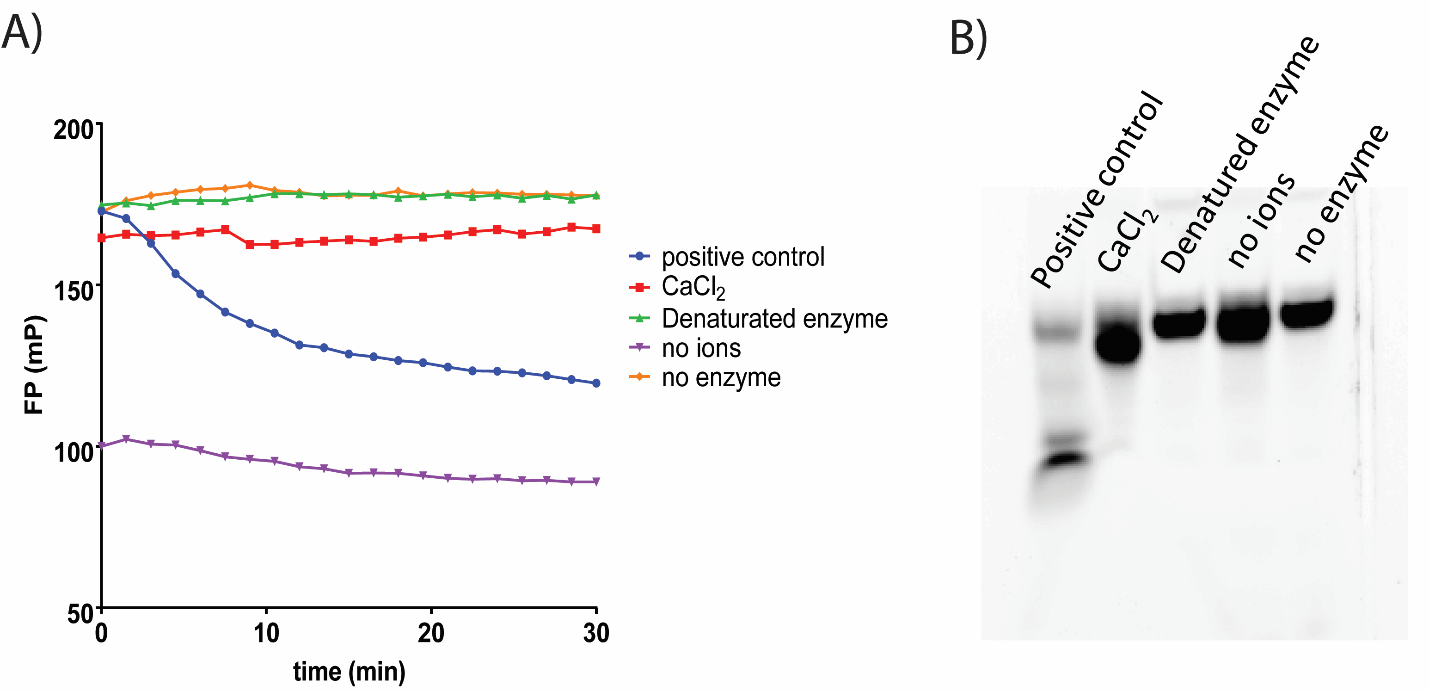

***Supplementary figure S3.*** Z' factor value for the high throughput screening assay. A) Scatter plot showing the 48 negative controls, the reaction mixture without ExoN and without inhibitor, represented as red dots, and 48 positive controls, the reaction mixture with ExoN and without inhibitor, represented as blue squares, for the library screening with MACV ExoN. The dashed lines show 3 SD for the negative controls (red dashed line) and 3 SD for the positive controls (blue dashed line). B) FP signal curves of the positive and negative controls used to calculate the Z’ value of MACV ExoN screening.

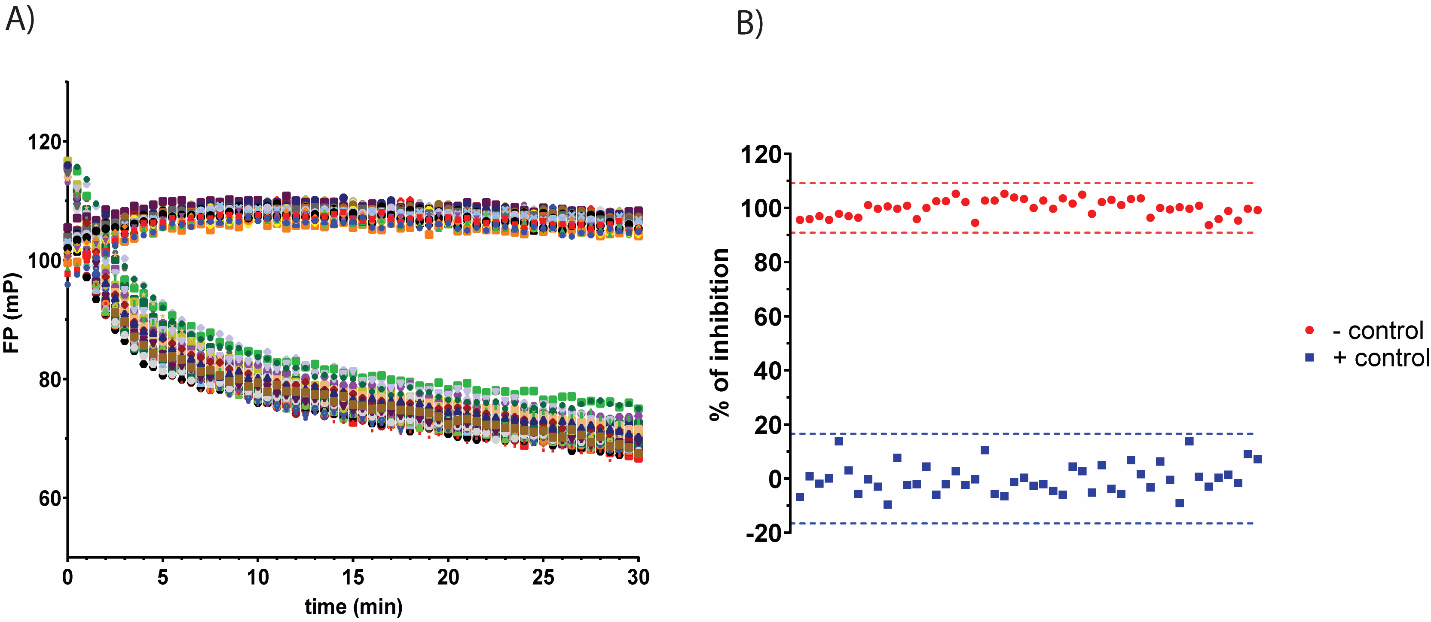

***Supplementary figure S4.*** Representative data for screening of inhibitors in SARS-CoV-2. A) FP curves of negative and positive controls. B) FP curves for the 23 ligands tested at 20 µM with the SARS-CoV-2 ExoN under conditions described in material and methods.

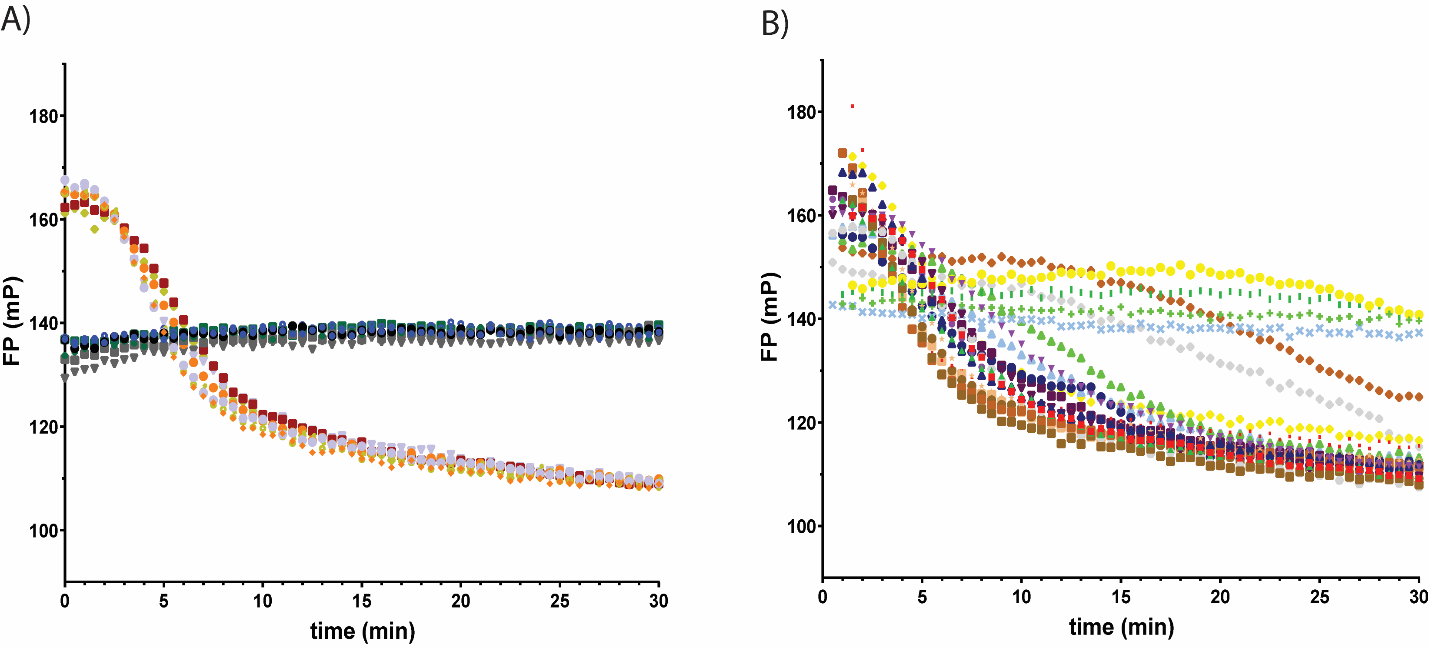

***Supplementary figure S5***. Screening of the focused library of metal chelators (23) against LCMV (panel A), MACV (panel B) and SARS-CoV-2 nsp14:nsp10 complex (panel C) ExoN followed by FP. The bars show the % of inhibition of the ExoN activity as described in materials and methods. For the screening conditions 100 nM 5'-FAM-RNA, 400 nM ExoN and 5 µM of inhibitor in 5% DMSO were used.

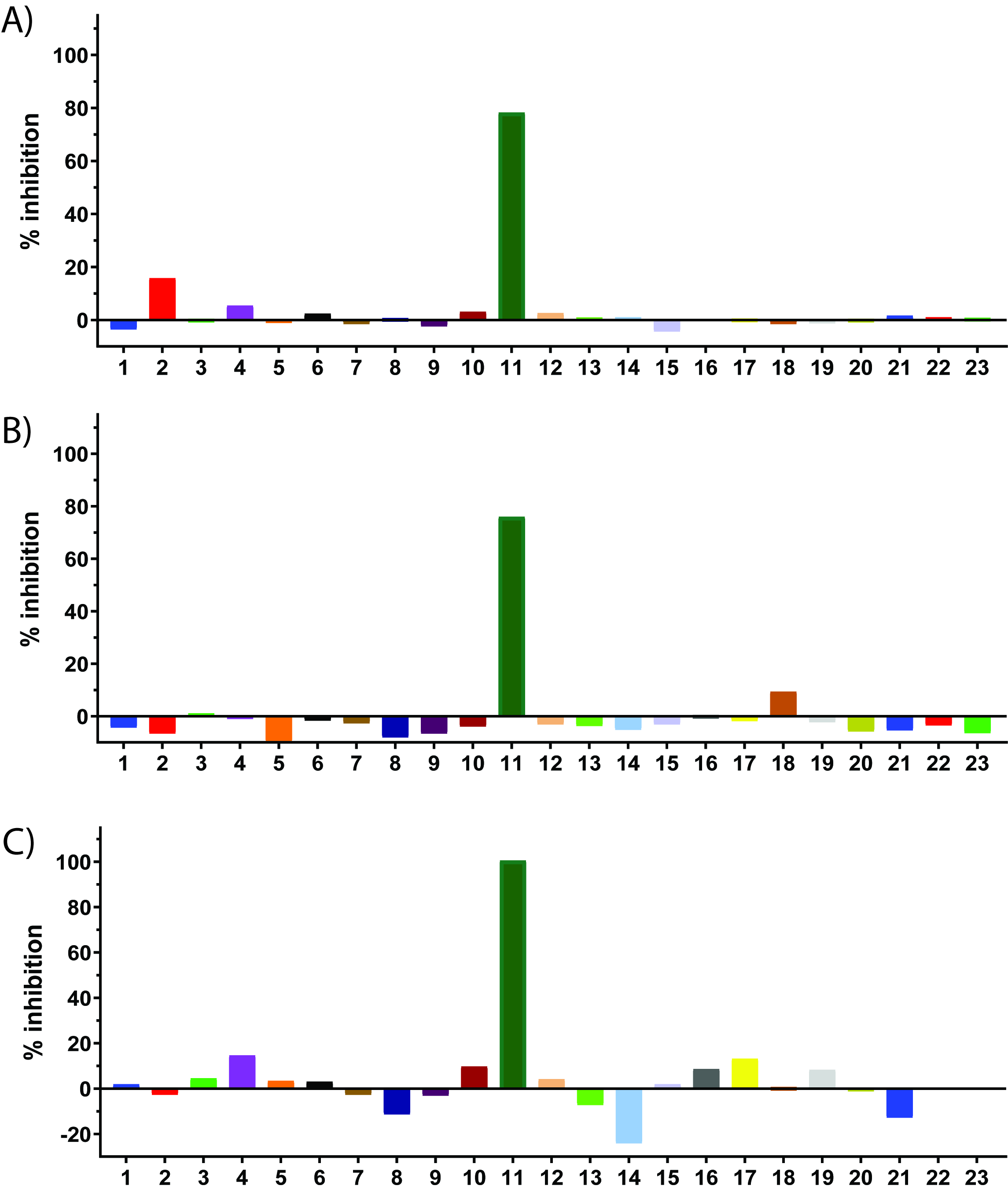
